# Highly heterogeneous long-range interactions explain chromosome compartmentalization

**DOI:** 10.1101/2019.12.29.890459

**Authors:** Y.A. Eidelman, I.V. Salnikov, S.V. Slanina, S.G. Andreev

## Abstract

The explanation for the compartmentalization of the chromosome contact maps observed by Hi-C method is still an unsolved mystery. The most natural and generally accepted explanation is that blocks of frequent/rare contacts on the entire map are associated with the existence of blocks of homogeneous elements along the chromosome that interact in 3D so that elements of the same type interact frequently and those of different types rarely. We study the polymer model of the chromosome, in which there are neither blocks of homogeneous elements nor homogeneous interactions and all the interaction energies are different. We demonstrate that such a heteropolymer model is able to describe chromosomal maps with high accuracy. The differences from compartment polymer models of chromosomes, which actually reflect the mechanism of microphase separation in polymers, are discussed further.

## Introduction

The study of chromosome structure compartmentalization is important. According to some data, compartmentalization correlates with gene expression, while according to other it does not. It changes in the course of differentiation and malignant transformation. In [1], the compartmentalization of contact maps obtained by the Hi-C method was explained by the physically natural and simplest assumption, as to the visible contact blocks being due to interaction of similar type elements located along the chromosomes. The number of element types proposed was two [1] or six [2], The question is, are there any other physical explanations?

In [3,4] we introduced a completely heteropolymer model of chromosomes, where all potentials between any polymer chain elements are different. The calculations with 100 kilo base pairs (kbp) resolution showed quite good ability of the model to capture the main properties of contact maps, observed in Hi-C experiments. What will change with increasing model resolution? Here we investigate the properties of the previous heteropolymer chromosome model by new calculations with higher resolutions, 50 and 10 kbp.

The distinct feature of model [3] is that it does not suggest that the genome/chromosome is partitioned into two [1] or six [2] types of compartments or subcompartments equivalent to two or six types of blocks of monomers. In [3,4] and in the present paper we explore the heterogeneous polymer where all subunits are assumed to be different and interaction energies between any pair of elements are different. To investigate the model’s ability to describe chromosome contact Hi-C data we calculate whole-chromosome contact maps with 50 and 10 kb resolution and compare the quality of description of experimental maps with respect to a polymer model with two to six types of chromatin subunits, or subcompartments, based on the idea of microphase separation. The conclusion we arrive at is that explanation of macroscopic compartmentalization first observed in [1] does not require a microphase separation mechanism as it is widely postulated.

## Results

### Heterogeneous polymer chromosome model describes the pattern of intrachromosomal contacts better than model postulating microphase separation

To test the descriptive capacity of the heterogeneous chromosome model, we used contact maps of chromosome 22 in GM12878 cells at higher resolution than previously used [3]. Fig.1 shows the results of comparison between simulated and experimental [2] contact maps with different resolutions.

**Fig.1.**
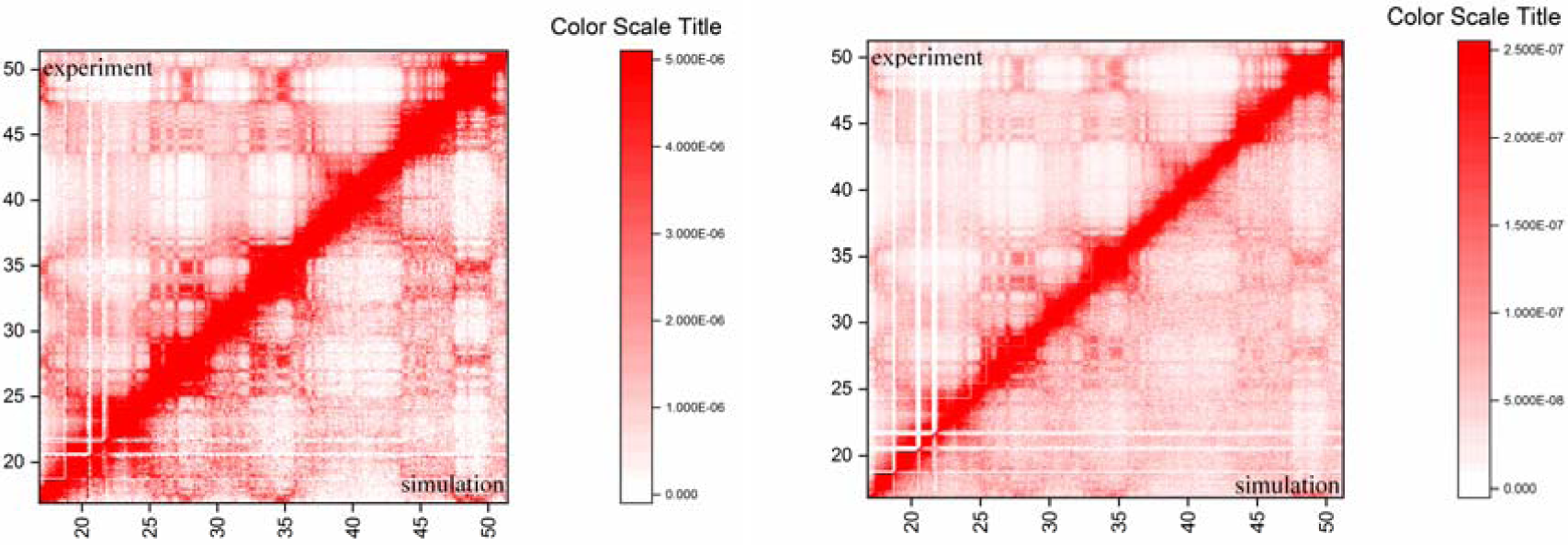
Heterogeneous polymer model describes Hi-C maps for chromosome 22 in GM12878 cells. Model resolutions: **A** – 50 kbp, **B** – 10 kbp. Experiment: [2], Pearson’s correlation: for 50 kbp R=0.982, for 10 kbp R=0.978.

Somewhat lower Pearson’s R and worse visual agreement for 10 kbp resolution calculations seen in Fig.1 is likely due to smaller statistics than required. This should be tested in the future.

For comparison, the result of calculations on the 6-subcompartment model with microphase separation [5] at 50 kbp resolution is presented in Fig.2.

**Fig.2.**
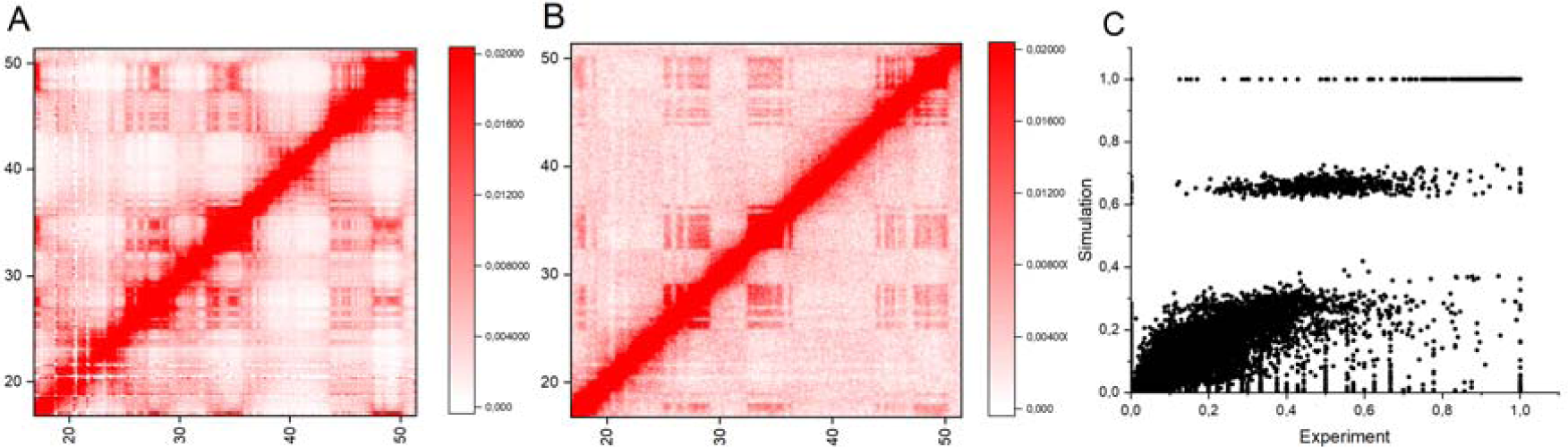
Simulation of chromosome 22 contact map by 6-subcompartment model is compared to Hi-C data [2], The model and its parameters were taken from [5]. Resolution 50 kbp.

### Checking the contact map compartments for uniformity

The concept of compartments imposes a number of conditions which should be met within the compartments: homogeneity of contacts, homogeneous potentials of interactions of the same type elements within the compartment of given type [1,2,5].

We checked these postulates by simulations with the most general, heteropolymer model, which describes the chromosome contact frequencies without assuming blocks of elements of the same type. To do this, we carried out A/B compartment analysis on the contact map according to [1], picked 15 map regions with boundaries corresponding to established A/B compartment borders (Fig.3) and tested them for the requirement of "homogeneity". All subunit pairs within each region belong to the same compartments (either A-A, or A-B, or B-B) and according to compartment concept, they should have similar contact frequencies. To offset the contact heterogeneity associated with the influence of genomic separation we used "observed/expected" map instead of raw contact map.

**Fig.3.**
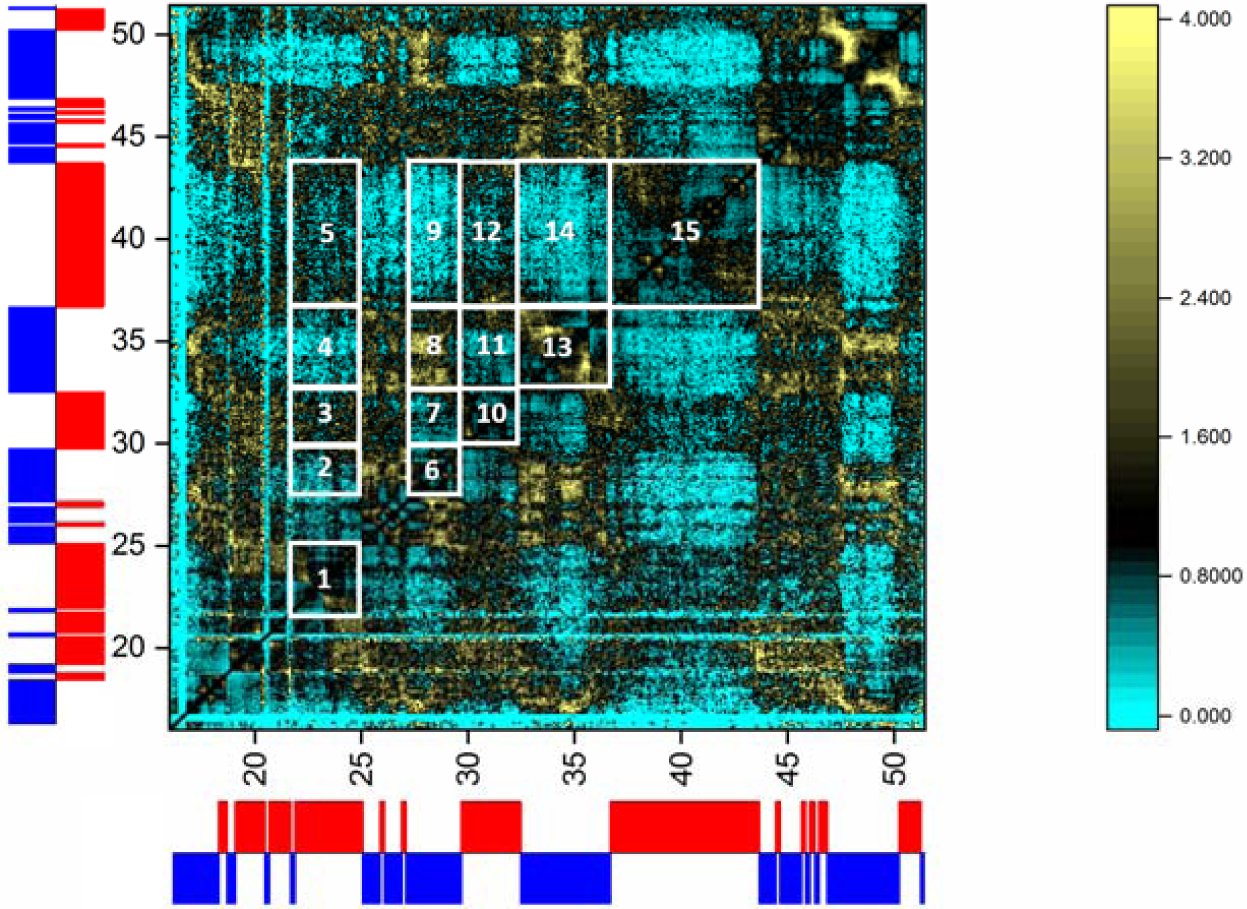
The test of simulated contact map (observed/expected) of chromosome 22 for contact heterogeneity analysis. The map was simulated by heterogeneous chromosome model, resolution 50 kbp, and coarsened to 100 kbp. To the left and below: ideogram of A/B compartments obtained by eigen-analysis according to [1] with 100 kbp resolution. A: red, B: blue. 15 regions picked for analysis together with their numbers are shown in the map (white).

One can conclude from Figures 3 and 4 that there are both heterogeneous distributions of contacts within compartment regions and strong deviations from the average values of frequency of contacts already at 50 kbp resolution. Moreover, the frequency of contacts of different types (A-B) does not differ much from the frequency of contacts of the same type (A-A or B-B), as should be expected from A-B compartment concept [1]. This fact directly contradicts the postulate of two compartments on the contact map of chromosome 22 with 50 kbp resolution and supports the heteropolymer model studied here.

**Fig.4.**
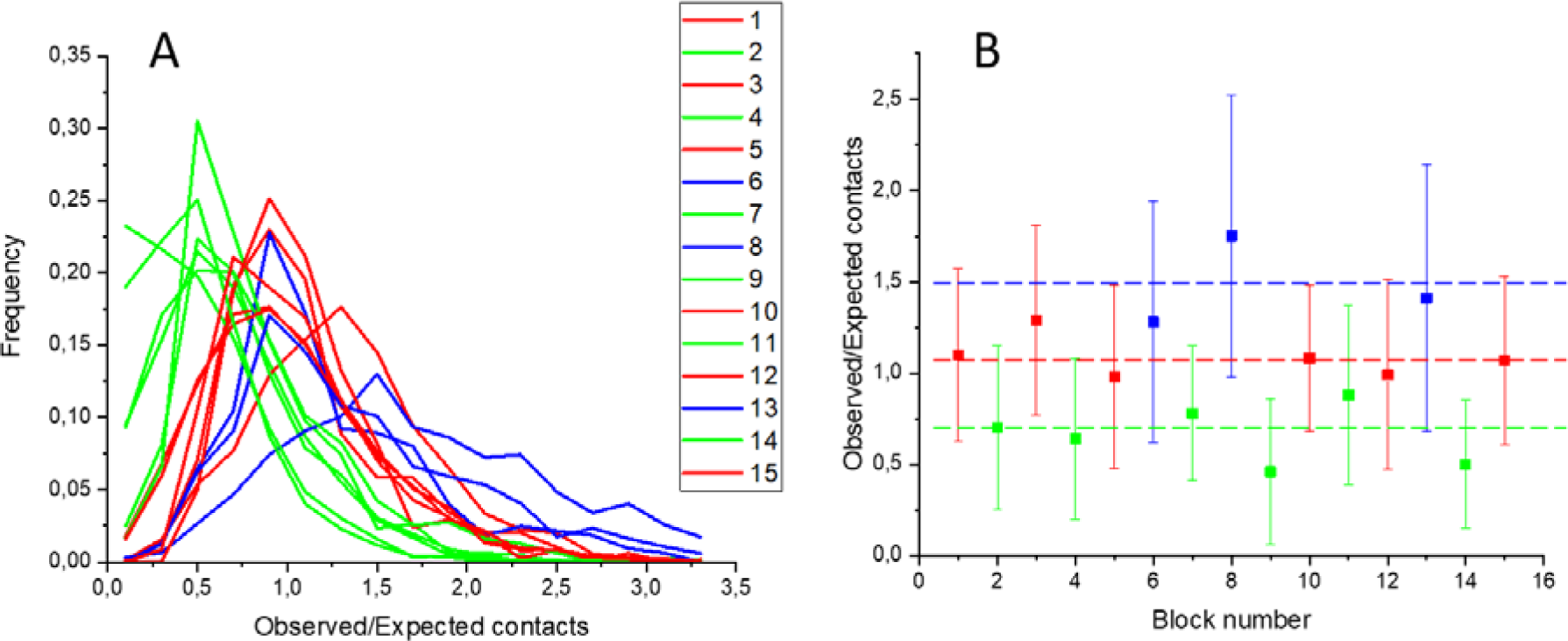
Analysis of contact heterogeneity in selected regions of chromosome 22 contact map with resolution 50 kbp. Region boundaries were determined according to borders of A/B compartments. **A**: distribution of contacts within each region. Region numeration is shown in Fig.3. Colors: red - A-A regions (i.e. both segments of the chromosome belong to A compartment, see Fig.3); blue - B-B regions; green - A-B regions. **B**: Mean number of contacts and standard deviation for each region. Dashed lines denote means for all regions of corresponding types.

Figures 5 and 6 show the results of the same analysis performed on maps with higher resolution, 10 kbp. The conclusion is the same as for 50 kbp resolution. Moreover, the heterogeneity of contact frequencies within each region is even higher and difference between same-type and different-type regions is even lower than for 50 kbp resolution which makes the compartment concept meaningless.

**Fig.5.**
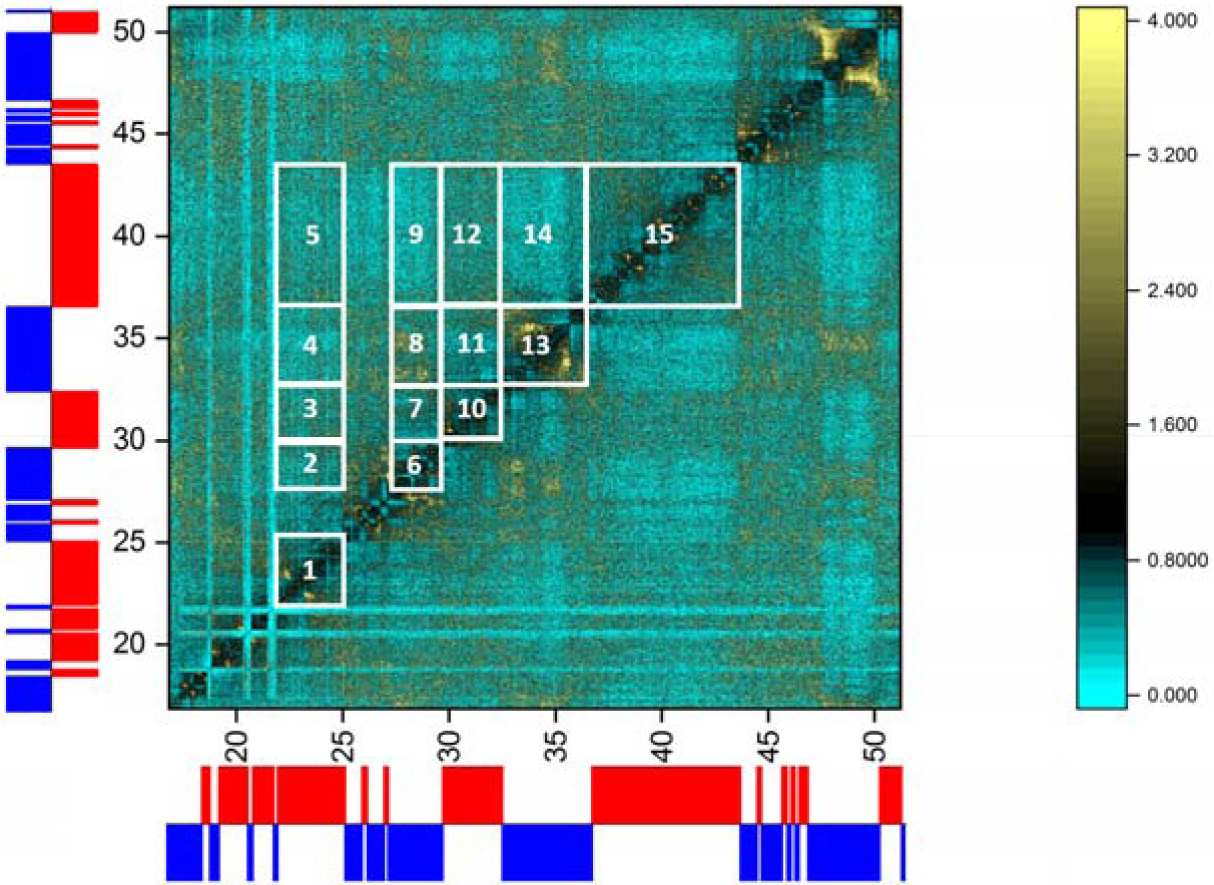
The simulated contact map (observed/expected) of chromosome 22 for contact heterogeneity analysis. Resolution 10 kbp. To the left and below: A/B compartments obtained with 100 kbp resolution. **A**: red, **B**: blue. The same 15 regions as for 50 kbp model (Fig.3) were picked for analysis (white).

**Fig.6.**
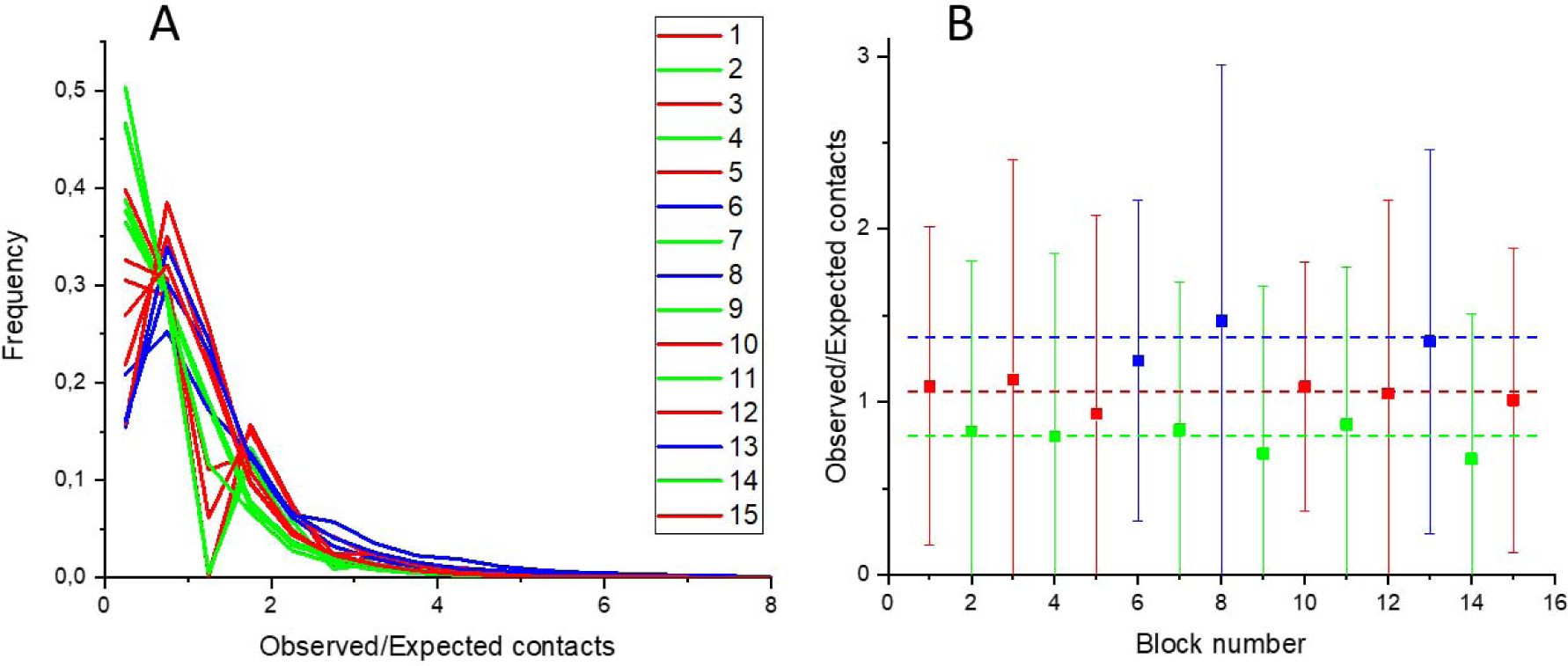
Analysis of contact heterogeneity in selected regions of chromosome 22 contact map with resolution 10 kbp. Region boundaries were determined according to borders of A/B compartments. **A**: distribution of contacts within each region. Region numeration is shown in Fig.5. Colors: red - A-A regions; blue - B-B regions; green - A-B regions. **B**: Mean number of contacts and standard deviation for each region. Dashed lines denote means for all regions of corresponding types.

The same analysis was repeated for 6-subcompartment concept [2] which was supposed to be more relevant than 2-compartment concept at resolutions as high as 50-100 kbp [2,5]. The results are presented in Figures 7-9. Only one type of distributions was visually different from others (A1-B2, resolution 50 kbp, Fig.8), but in a whole picture the previous conclusion stands: heterogeneity within each region is higher than between same-type and different-type regions. This contradicts the postulate of six subcompartments on the contact map of chromosome 22 with both 50 and 10 kbp resolution and supports the proposed heteropolymer model.

**Fig.7.**
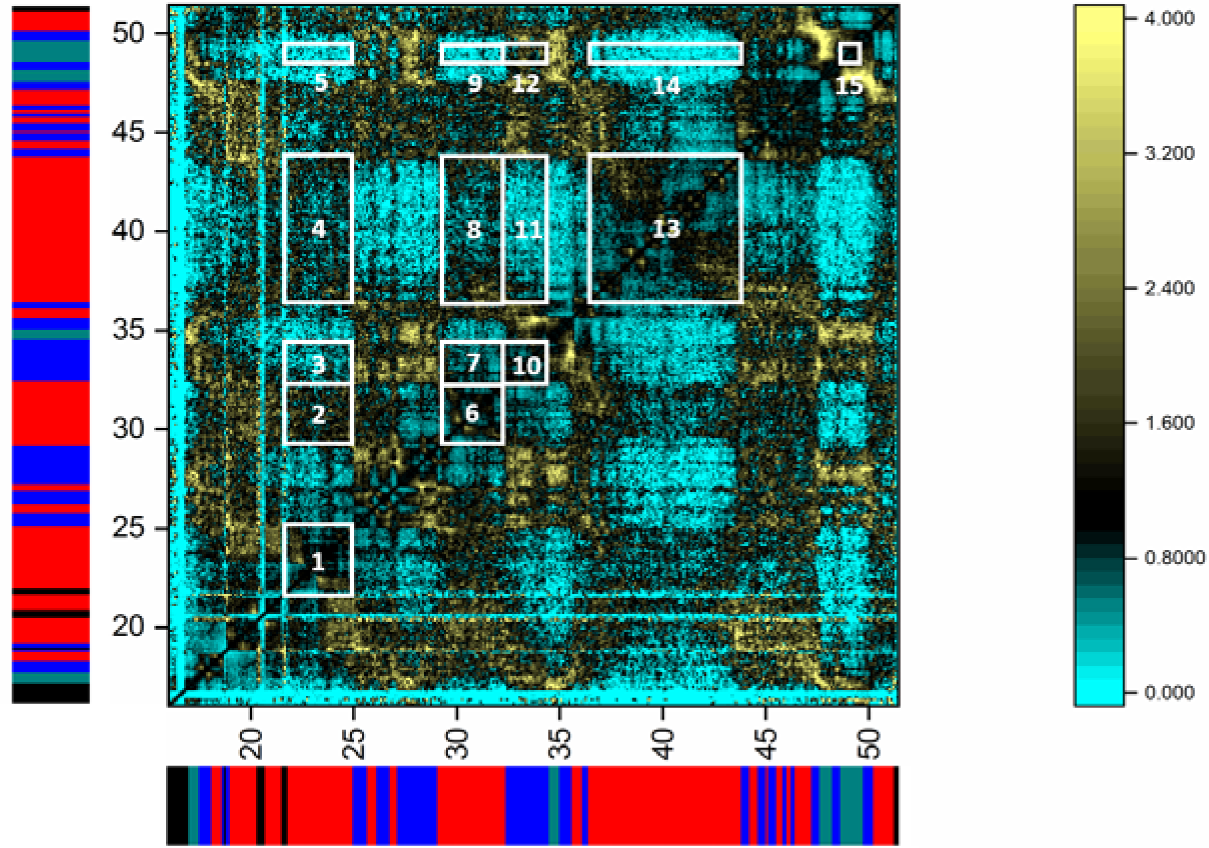
The simulated contact map (observed/expected) of chromosome 22 for contact heterogeneity analysis. Heterogeneous chromosome model, resolution 50 kbp, map coarsened to 100 kbp. To the left and below: ideogram of A1/B1/B2 subcompartments from [2], 100 kbp resolution. A1: red, B1: blue, B2: teal, N/A: black. A2 and B3 subcompartments are not present in chromosome 22. The map was divided into regions according to subcompartment boundaries. 15 regions picked for analysis together with their numbers are shown in the map (white). Their positions are partially different from those picked previously (Figs 3, 5) as subcompartment boundaries are different from A/B case.

**Fig.8.**
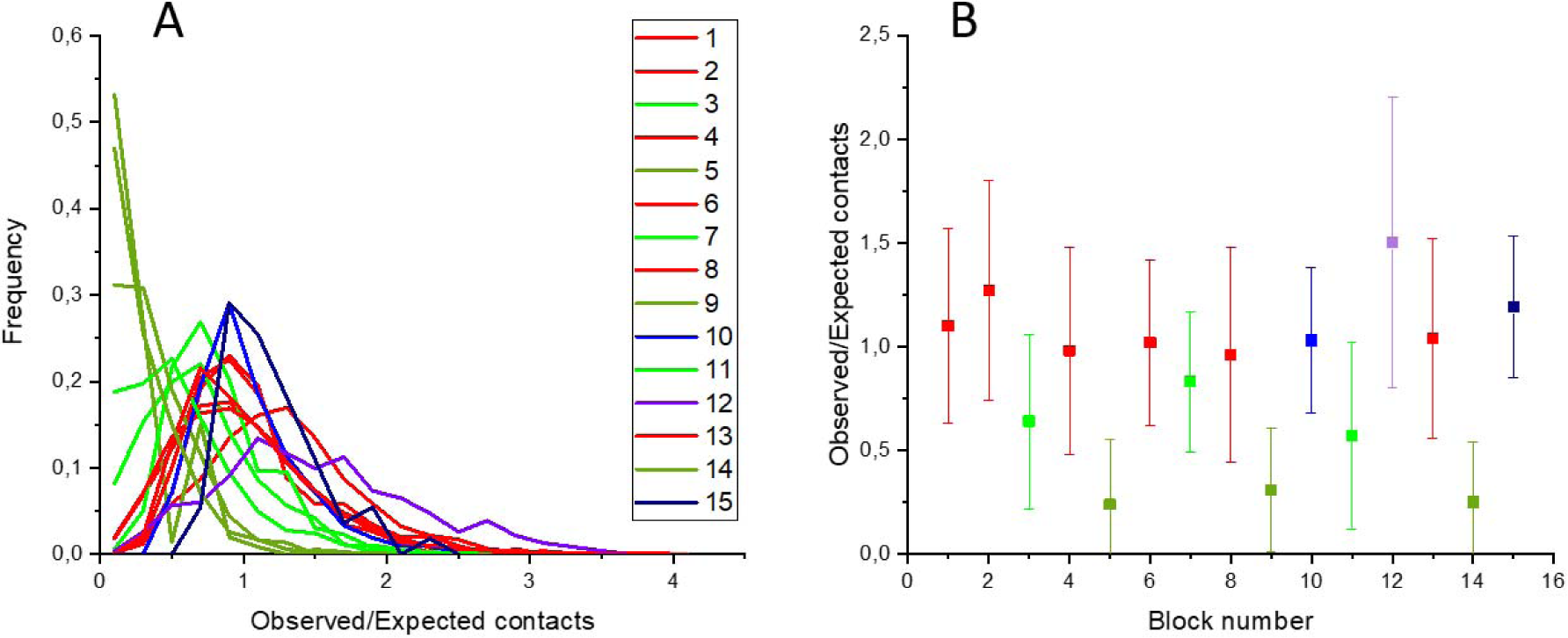
Analysis of contact heterogeneity in selected regions of chromosome 22 contact map with resolution 50 kbp. Region boundaries were determined according to borders of A1/B1/B2 subcompartments. **A:** distribution of contacts within each region. Region numeration is shown in Fig.7. Colors: red - Al-Al regions; blue - Bl-Bl regions; dark blue - B2-B2 regions; green - Al-Bl regions; dark green - A1-B2 regions; violet - B1-B2 regions. **B:** Mean number of contacts and standard deviation for each region.

**Fig.9.**
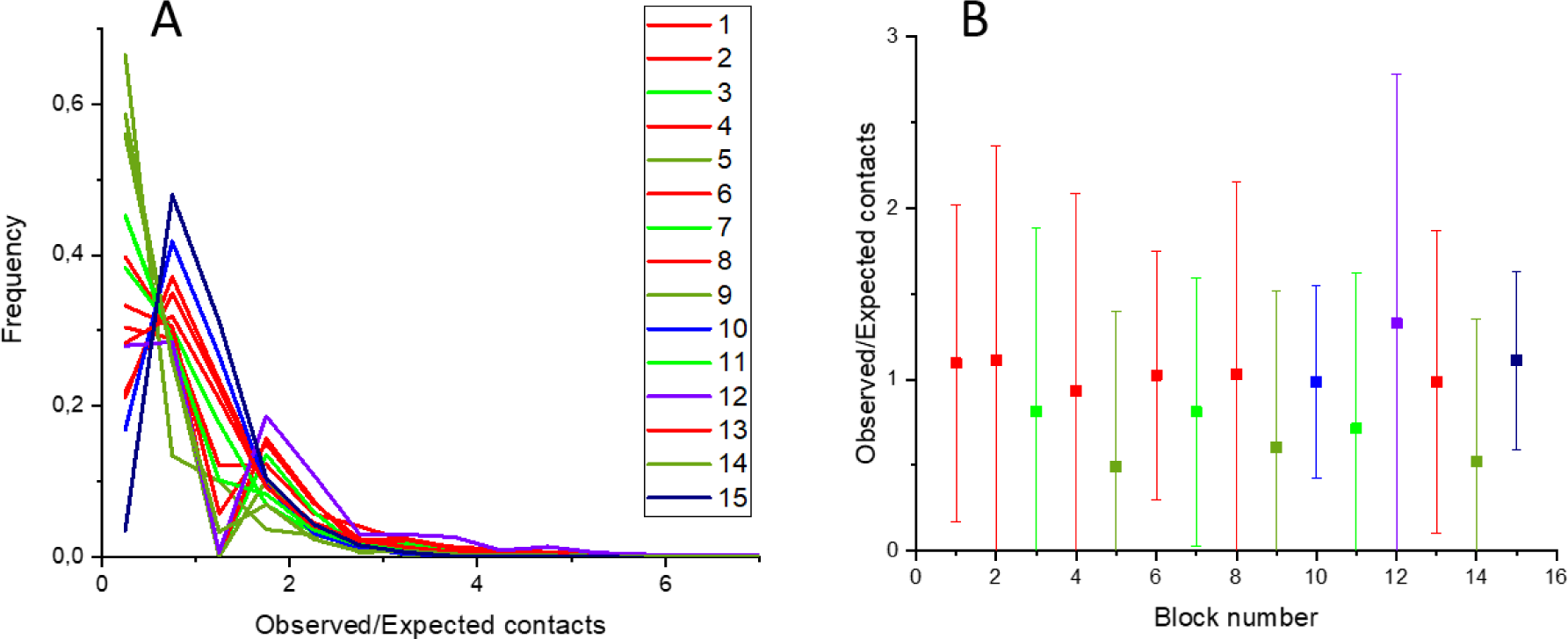
Analysis of contact heterogeneity in selected regions of chromosome 22 contact map with resolution 10 kbp. Region boundaries were determined according to borders of A1/B1/B2 subcompartments. **A**: distribution of contacts within each region. Region positions and numeration and the same as for 50 kbp resolution (Figures 7, 8). Colors: red - Al-Al regions; blue - Bl-Bl regions; dark blue - B2-B2 regions; green - Al-Bl regions; dark green - A1-B2 regions; violet - B1-B2 regions. **B**: Mean number of contacts and standard deviation for each region.

Figures 10-12 show the presence of wide potential distributions (heterogeneity) necessary to describe the contact frequencies in different regions with 50 and 10 kbp resolutions according to our heterogeneous model. They are very different from the homogeneous potentials in regions, which are postulated by a model with a microphase separation at 50 kbp resolution [5]. For 10 kbp the potential spread is even wider than for 50 kbp, which indicates the heterogeneity of interactions and does not support the assumption of homogeneous blocks of similar elements and, accordingly, blocks of identical interaction energies between repeating elements [1,2,5].

**Fig.10.**
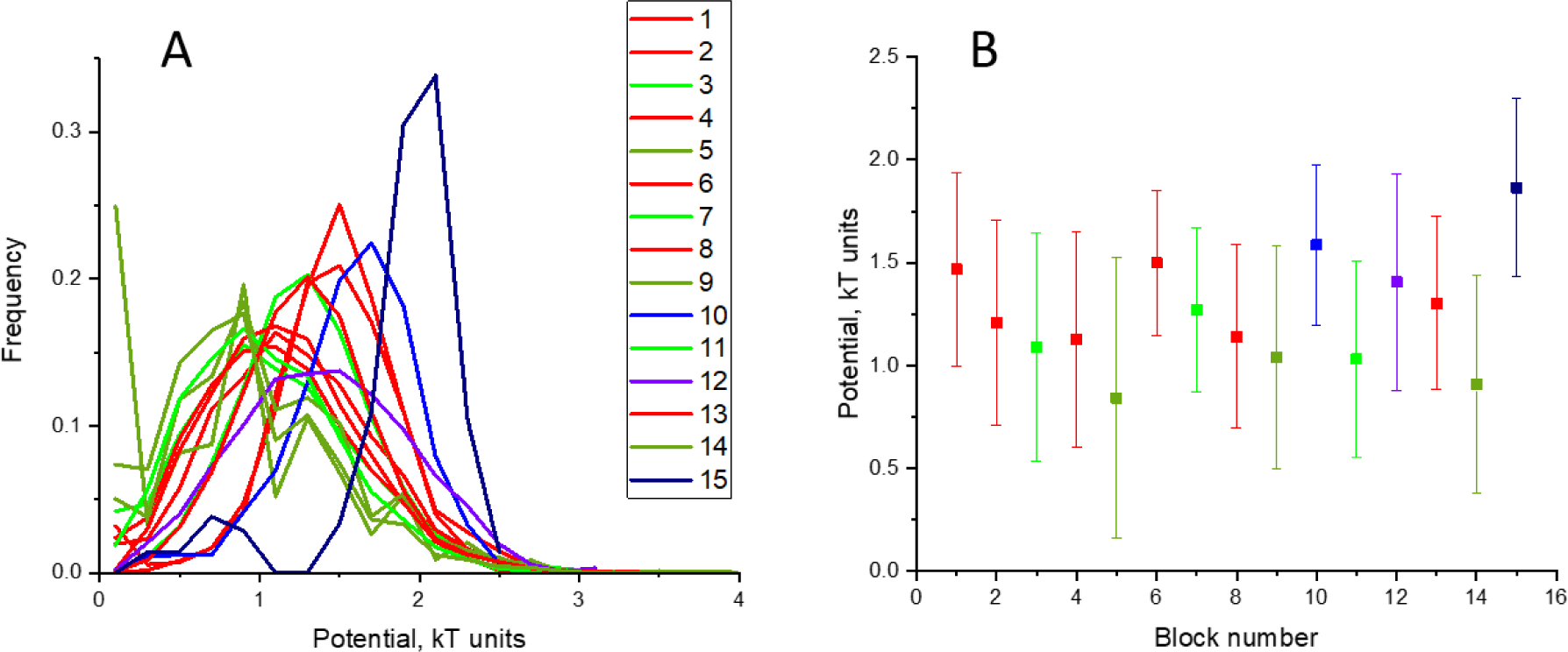
Analysis of potential heterogeneity in selected regions of chromosome 22 contact map with resolution 50 kbp. Regions are the same as in contact heterogeneity analysis (Fig.8). **A**: distribution of potentials within each region. Region numeration is shown in Fig.7. Colors: red - Al-Al regions; blue - Bl-Bl regions; dark blue - B2-B2 regions; green - Al-Bl regions; dark green - A1-B2 regions; violet - B1- B2 regions. **B**: Mean potential and standard deviation for each region.

**Fig.11.**
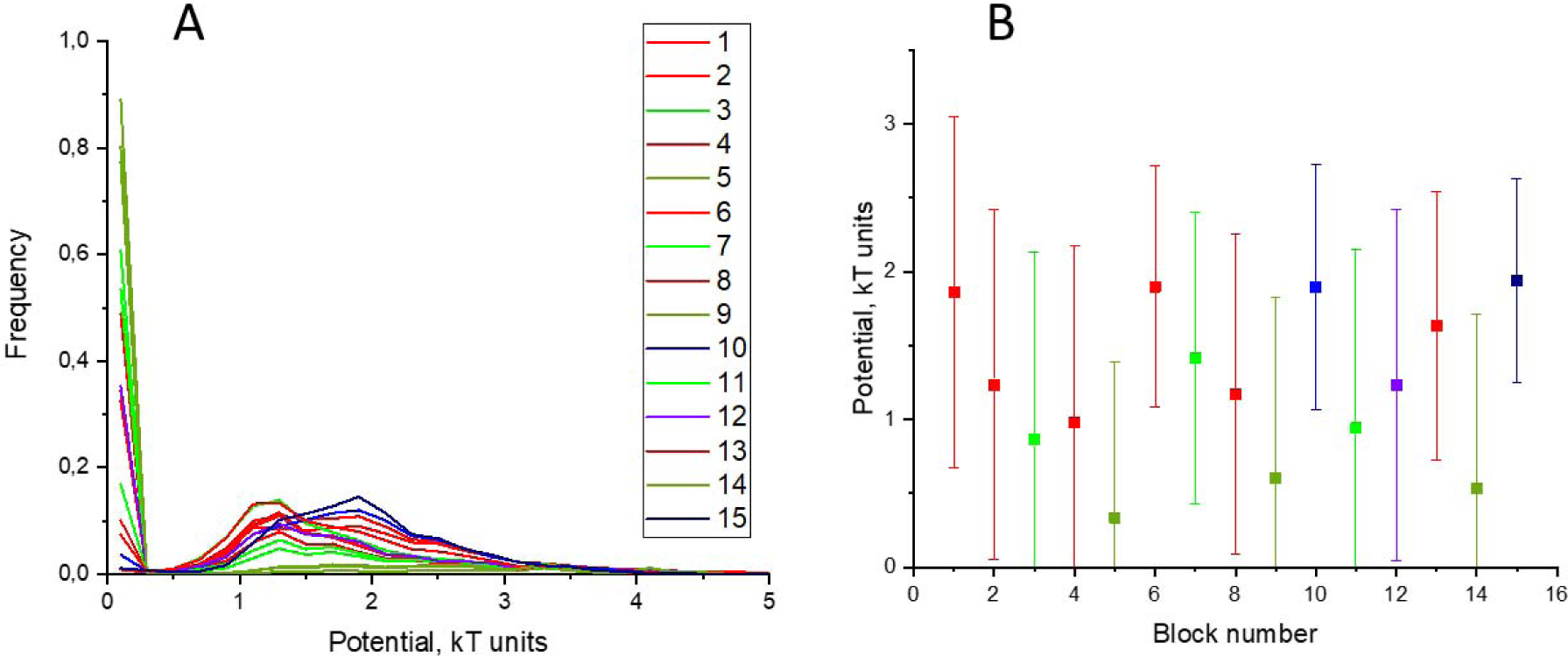
Analysis of potential heterogeneity in selected regions of chromosome 22 contact map with resolution 10 kbp. Regions are the same as in contact heterogeneity analysis (Fig.8). **A**: distribution of potentials within each region. Region numeration is shown in Fig.7. Colors: red - Al-Al regions; blue - Bl-Bl regions; dark blue - B2-B2 regions; green - Al-Bl regions; dark green - A1-B2 regions; violet - B1- B2 regions. **B**: Mean potential and standard deviation for each region.

**Fig.12.**
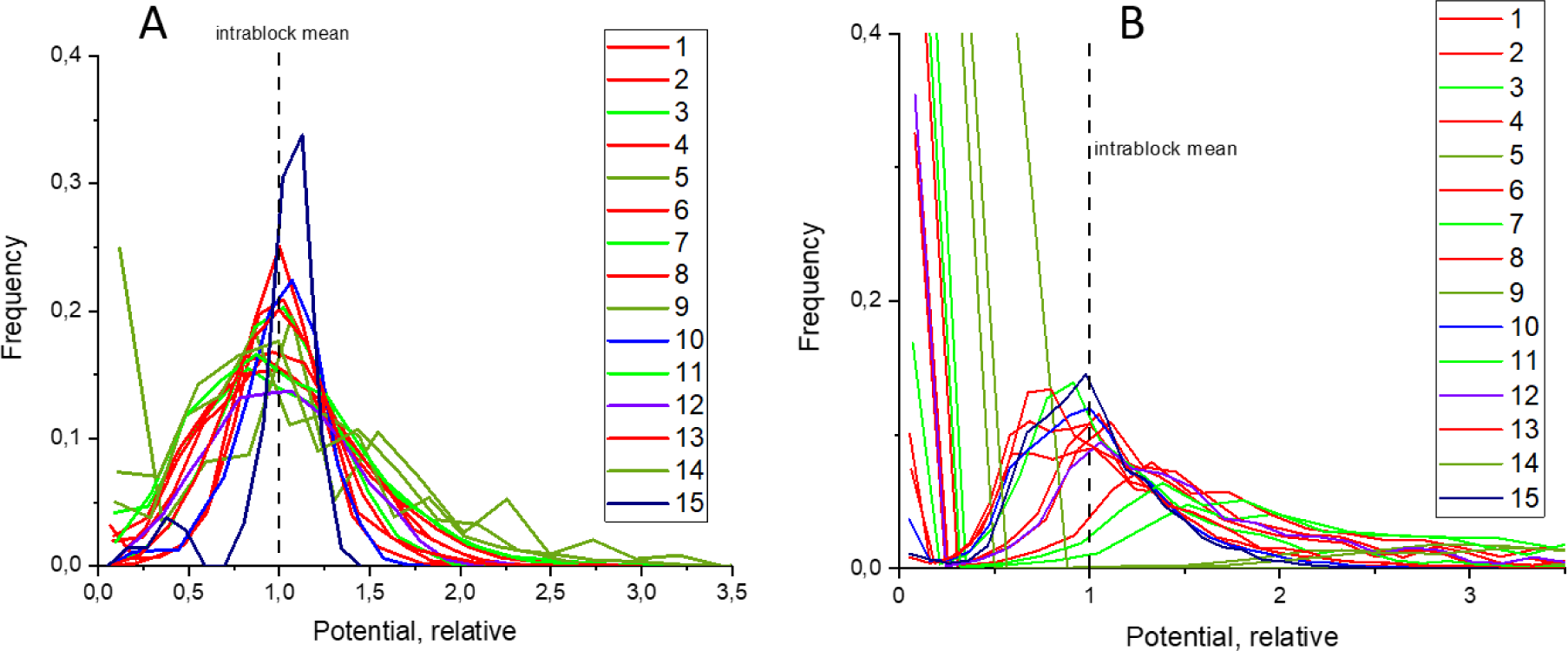
Potentials within each region are highly heterogeneous. A: distribution for relative potentials for 50 kbp simulation. B: for 10 kbp simulation. Relative potentials were obtained for each region by dividing potentials by the mean for this region. Thus, all means equal unity (vertical dashed lines). Region numbers and line colors are the same as in corresponding distributions for absolute potentials (Figures 10, 11).

Fig.13 demonstrates that heterogeneous contacts are more adequately described by heterogeneous potentials that homogeneous one for the given region or compartment.

**Fig.13.**
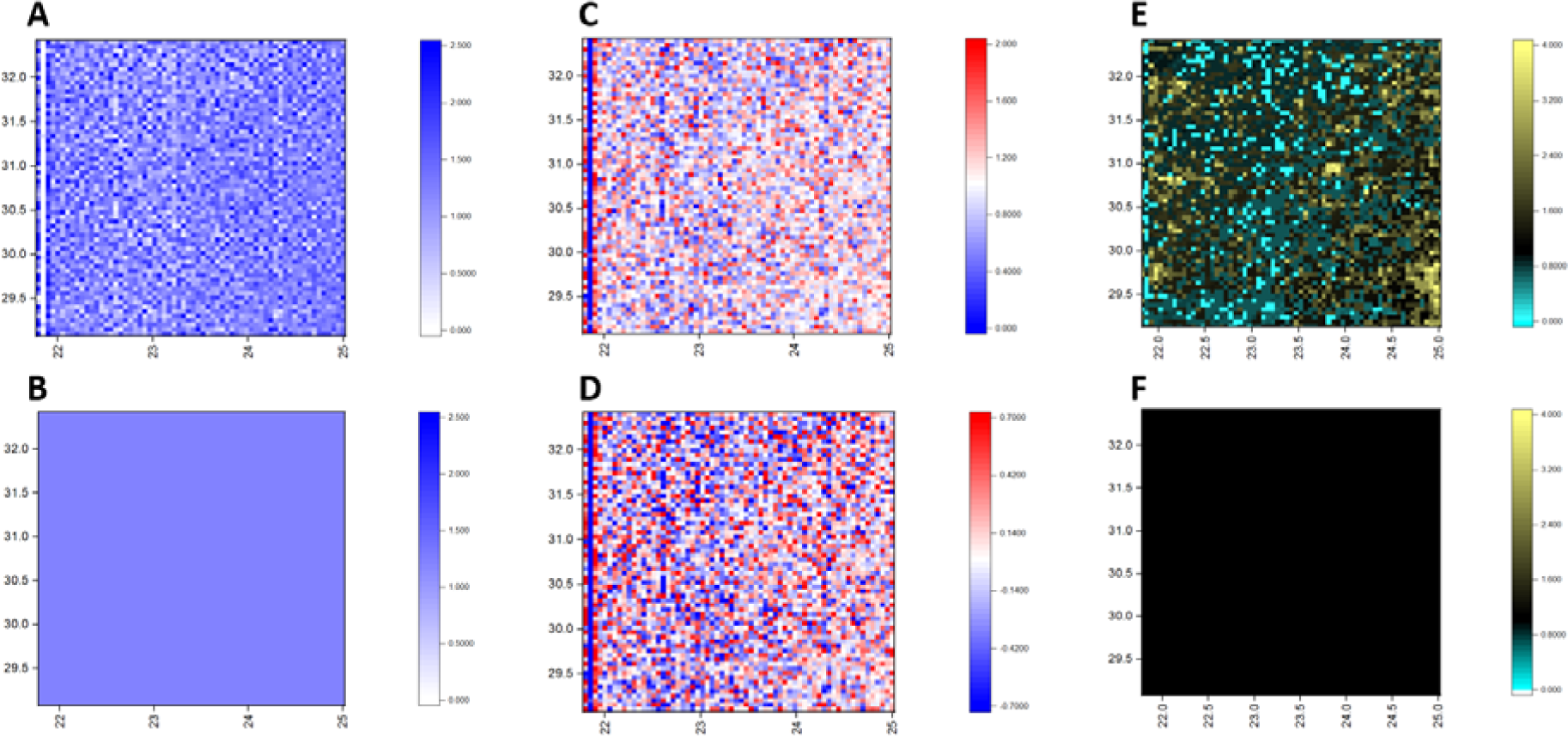
Interconnection between heterogeneity of potentials, i.e. absence of (sub)compartments, and heterogeneity of contacts. As an example, region 2 for 50 kbp resolution model is taken, 21.8-25 × 29.1-32.4 Mbp, A1-A1 type (see Fig.7). **A**: simulated heterogeneous map of potentials. **B**: the corresponding map for the case with all potentials equal to the mean (<u>=1.21 kT). **C** and **D** illustrate potential heterogeneity. **C**: simulated map of relative potentials u_1_(i,j)=u(i,j)/<u>. **D**: simulated map of potential relative deviation u_2_(i,j)=(u(i,j)-<u>)/<u>. **E**: contact map (observed/expected) obtained with heterogeneous potentials in panel A. **F**: contact map assumed from homogeneous potentials (panel B).

Fig.14 shows the difference in interaction energy along the chromosome between the heteropolymer chromosome model and the model based on microphase separation.

**Fig.14.**
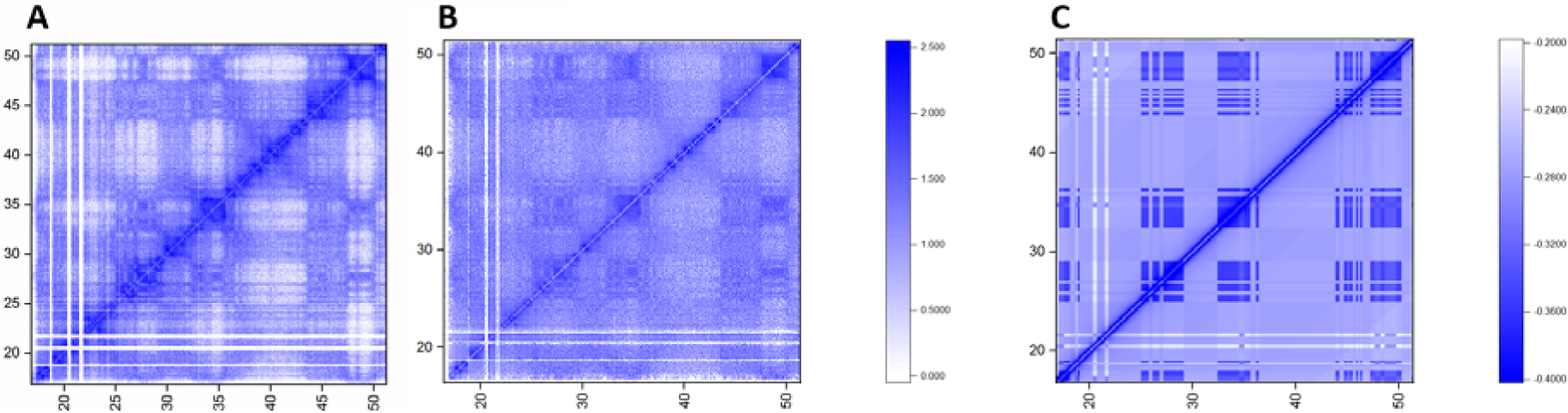
Maps of potentials for different models of chromosome 22. **A**: heterogeneous model, resolution 10 kbp. **B**: heterogeneous model, resolution 50 kbp. C: 5-subcompartment model [5], resolution 50 kbp. The values are in kT units.

## Discussion

We conclude that there are heterogeneous distributions of contact densities in different parts of chromosome which are due to heterogeneous interaction energies. Chromosome compartmentalization can be explained not by microphase separation of monomer blocks along the chromosome but by effect of heterogeneous network of volumetric interaction potentials in chromosome. A consequence of this approach is that there are sequence-specific (i.e. depending on position of each interacting elements) heterogeneous volumetric potentials which represent on molecular level the variety of crosslinking factors, (multi)protein complexes, transcription factors, ncRNA, *etc*. They can promote specificity of interaction energies in chromosome on different scales. The macro-compartmentalization seen on low-resolution maps may be partly the result of map coarsening. At high resolution there are wide distributions in all parameters (contacts, potentials) in all blocks which, when resolution is decreased and the map is coarsened, produce the effect of distinct blocks, sharp borders, compartments, etc. Homogeneous blocks of same-type elements at the microscopic level are not observed after analyzing the contact map with high resolution (50 and 10 kbp). Alternatively, we cannot rule out that the frequency jumps and skips in the maps and the resulting heterogeneity of interaction energies may be related to not very high-quality maps, and not to the heterogeneous nature of long-range interactions in chromosomes.

In conclusion, the main result of the introduction of a new heteropolymer model of the chromosome is as follows. Explanation of the compartmentalization of the contact map is possible without the mechanism of microphase separation. In other words, contact maps can be explained without blocks of same-type elements along the chromosome. Our model does not require chromatin subcompartments along the chromosome to understand chromosome compartmentalization observed by proximity ligation technique [1]. Although the mechanism of microphase separation is physically simplest and natural for the explanation of the checkerboard type of contact maps, it is not the only one and, as we show, not necessary.

## Methods

The interphase structure of human chromosome 22 was modeled with the inhouse software using molecular dynamics package OpenMM. A chromosome was modeled as a heteropolymer chain of subunits with either 50 kbp or 10 kbp DNA content. The interaction potential between any pair of non-neighbor subunits (/’,;’) consists of excluded volume potential and attraction potential. The former is the same for all pairs of subunits, while the latter depends on the position of the subunits along the chain. This is similar to the method developed by us in [2,3].

The coefficients for the attraction potential are determined to provide the best fit of the experimental Hi-C maps by the model. To this end we developed the iterative algorithm described in [2,3].

## Acknowledgements

The study was supported by MEPhI Academic Excellence Project (Contract No. 02.a03.21.0005); Russian Foundation for Basic Research grant 14-01-00825; The research is carried out using the equipment of the shared research facilities of HPC computing resources at Lomonosov Moscow State University and Joint Supercomputer Center of the Russian Academy of Sciences.

## Author Contributions

Y.E.: software, scientific analysis, work on manuscript. I.S.: software. S.S.: data acquisition and analysis. S.A.: conceptualization, scientific analysis, work on manuscript.

